# LILRB1 and LILRB2 expression in peripheral blood immune cells at 18 and 24 months of age in infants born from mothers with placental malaria

**DOI:** 10.1101/2021.10.26.465873

**Authors:** Celia Dechavanne, Odilon Nouatin, Rafiou Adamou, Sofie Edslev, Anita Hansen, Florian Meurisse, Ibrahim Sadissou, Erasme Gbaguidi, Jacqueline Milet, Gilles Cottrell, Laure Gineau, Audrey Sabbagh, Achille Massougbodji, Kabirou Moutairou, Eduardo A. Donadi, Edgardo Carosella, Philippe Moreau, Ed Remarque, Michael Theisen, Nathalie Rouas-Freiss, André Garcia, Benoit Favier, David Courtin

## Abstract

**Background:** Placental malaria (PM) is associated with a higher susceptibility of infants to *Plasmodium falciparum (Pf)* malaria. A hypothesis of immune tolerance has been suggested but no clear explanation has been provided so far. Our goal was to investigate the involvement of inhibitory receptors LILRB1 and LILRB2, known to drive immune evasion upon ligation with pathogen and/or host ligands, in PM-induced immune tolerance.

**Methods:** Infants of mothers with or without PM were enrolled in Allada, southern Benin, and followed-up for 24 months. Antibodies with specificity for five blood stage parasite antigens were quantified by ELISA, and the frequency of immune cell subsets was quantified by flow cytometry. LILRB1 or LILRB2 expression was assessed on cells collected at 18 and 24 months of age.

**Results:** Infants born to PM-mothers had a higher risk of developing clinical malaria than those born to mothers without PM (IRR=1.53, p=0.040), and such infants displayed a lower frequency of non-classical monocytes (OR=0.74, p=0.01) that overexpressed LILRB2 (OR=1.36, p=0.002). Moreover, infants born to PM-mothers had lower levels of cytophilic IgG and higher levels of IL-10 during active infection.

**Conclusion:** Modulation of IgG and IL-10 levels could impair monocyte functions (opsonisation/phagocytosis) in infants born to PM-mothers, possibly contributing to their higher susceptibility to malaria. The long-lasting effect of PM on infants’ monocytes was notable, raising questions about the capacity of ligands such as Rifins or HLA-I molecules to bind to LILRB1 and LILRB2 and to modulate immune responses, and about the reprogramming of neonatal monocytes/macrophages.

**KEY POINTS:** Infants of mothers with placental malaria were more susceptible to clinical malaria than those born to mothers without placental malaria and they displayed a lower frequency of non-classical monocytes that overexpressed LILRB2.

## Introduction

Each year, up to 125 million pregnancies occur in malaria endemic countries, with a heightened risk of poor outcomes including miscarriage, maternal death and severe anaemia (1, 2). Pregnancy-associated malaria is also deleterious for the newborn (3), leading to low birth weight and increasing the risk of infant morbidity and mortality (1). Placental malaria (PM) due to *Plasmodium falciparum (Pf)* is estimated to cause up to 100,000 infant deaths every year (4). Children born to mothers with PM are more susceptible to malaria (3). PM shortens the delay to first malaria infection (5–9), and this is thought to be due to a phenomenon named immune tolerance (IT). This phenomenon may be driven by fetal sensitization to malaria antigens *in utero* leading to a modification of immune development of the foetus (10, 11). At present, no unequivocal explanation has been proposed.

Leukocyte immunoglobulin like receptor B (LILRB)1 and LILRB2 are inhibitory receptors that play an important role in the regulation of immune responses that modulate progression or control of infectious diseases (12, 13). LILRB2 is exclusively expressed by myeloid cells including monocytes, dendritic cells and neutrophils (12). In contrast, LILRB1 is found on monocytes and dendritic cells but also on B cells and subsets of CD8 T, γδ T and NK cells (12). Binding of LILRB1 and LILRB2 to HLA-I molecules affects the function of the corresponding immune cell populations, thereby modulating crucial steps in the immune response such as cell differentiation, migration, proliferation, cytotoxicity and cytokines or antibody production. Recent studies indicate a complex interplay between monocytes and malaria infection. Indeed, monocytes play an important role in the immune responses against malaria through phagocytosis and cytokine production. However, exacerbated activation of monocytes could also increase the level of inflammation, leading to detrimental host immune responses (14). The non-classical HLA class I molecule HLA-G, known to be involved in maternal maternofetal tolerance, presents the highest affinity for binding to LILRB1 and LILRB2. HLA-G and IL10 mutually up-regulate their expression, and are involved in neonatal immunoregulatory mechanisms (15). Of note, high plasma levels of HLA-G were previously associated with an increased risk of *Pf* infection in infants (16, 17).

In the present study, we aimed to define the role of PM on immune profiles as well as on the level of anti-malarial antibodies in infant with a particular focus on LILRB1, LILRB2, IL-10 and HLA-G expression. These data bring a better understanding of the dynamics of monocyte subsets and LILRB1 / LILRB2 inhibitory receptor expression during *Pf* infection with potential implications for the design of new therapeutic strategies against malaria.

## Methods

### Study design and follow-up

The present follow-up is part of a study concerning 1,183 pregnant women participating in *Malaria in Pregnancy Preventive Alternative Drugs*, a randomized trial of intermittent preventive treatment (IPTp) in Benin (18). The first 154 infants for which flow cytometry data were available were enrolled from January 2010 to June 2011 and followed throughout the first 2 years of life in the TOLIMMUNPAL project (19). At birth, newborn’s gender, weight and axillary temperature were recorded. At 6, 9, 12, 18 and 24 months, a medical questionnaire was filled-out. A thick blood smears (TBS) was monthly performed to detect asymptomatic malaria carriage. A rapid diagnosis test (RDT) for malaria was performed during the follow-up if an infant was febrile. A symptomatic malaria attack was defined as the presence of fever (or a 24 hours’ history of fever) and a positive RDT and/or a positive TBS. Malaria attacks were treated with artemether-lumefantrine combination, as recommended by the Beninese National Malaria Control Program. An asymptomatic infection was defined as a positive monthly systematic TBS with no fever or no history of fever. Peripheral blood at 18 and 24 months was collected to perform the cellular immune cell phenotyping including LILRB1 and LILRB2 expression, to determine antibodies, cytokines and HLA-G concentrations.

Because the environmental risk of exposure to malaria may play a role in infants’ susceptibility to malaria infection, mosquito catches were performed monthly over two consecutive nights, throughout the course of the study, to assess the density of malaria vectors. A time- and space-dependent environmental risk of exposure was assessed for each child by a predictive model (20).

### Antibody, cytokine and HLA-G measurements

The Enzyme Linked ImmunoSorbent Assay (ELISA) standard operating procedures developed by the African Malaria Network Trust was used to assess antibody concentrations to MSP1, MSP2, MSP3, AMA1 and GLURP antigens. All details about the recombinant proteins and the procedure are described by Dechavanne et al.(21).

Concentrations of IL-4, IL-10, TNF-α and IFN-γ were determined using BD OptEIA ELISA Set (BD Biosciences, San Diego, CA, USA) following the manufacturer’s protocol and as previously described by our team (22). The levels of soluble HLA-G isoforms HLA-G1 and HLA-G5 were determined by ELISA assay as we described previously (17).

### Immune cell phenotyping and LILRB1 and LILRB2 expression

Phenotype analysis was performed on infant peripheral blood (1 mL) the day of collect. The expression of LILRB1 and/or LILRB2 inhibitory receptors on the surface of the following cell populations was analyzed: T helper cells (CD3^+^CD4^+^), T cytotoxic cells (CD3^+^CD8^+^), T regulatory cells (CD3^+^CD4^+^CD25^high^CD127^-^), T effector cells (CD3^+^CD4^+^CD25^+^CD127^+^), γδ T cells (CD3^+^Vd2^+^) and B cells (CD19^+^). NK cell subsets were characterized by expression of CD56 and CD16 among the mononuclear lymphoid cells. Blood monocyte subpopulations were characterized by expression of CD14 and CD16 markers in the population of mononuclear cells. Neutrophils and eosinophils are respectively CD16^+^ and CD16^-^ in the polymorphonuclear cell population. To define positive and negative events, isotype-matched control antibodies were used. Flow cytometer data were analyzed using FlowJo software version 10. The gating strategy is presented in Figure 1.

**Figure 1.**
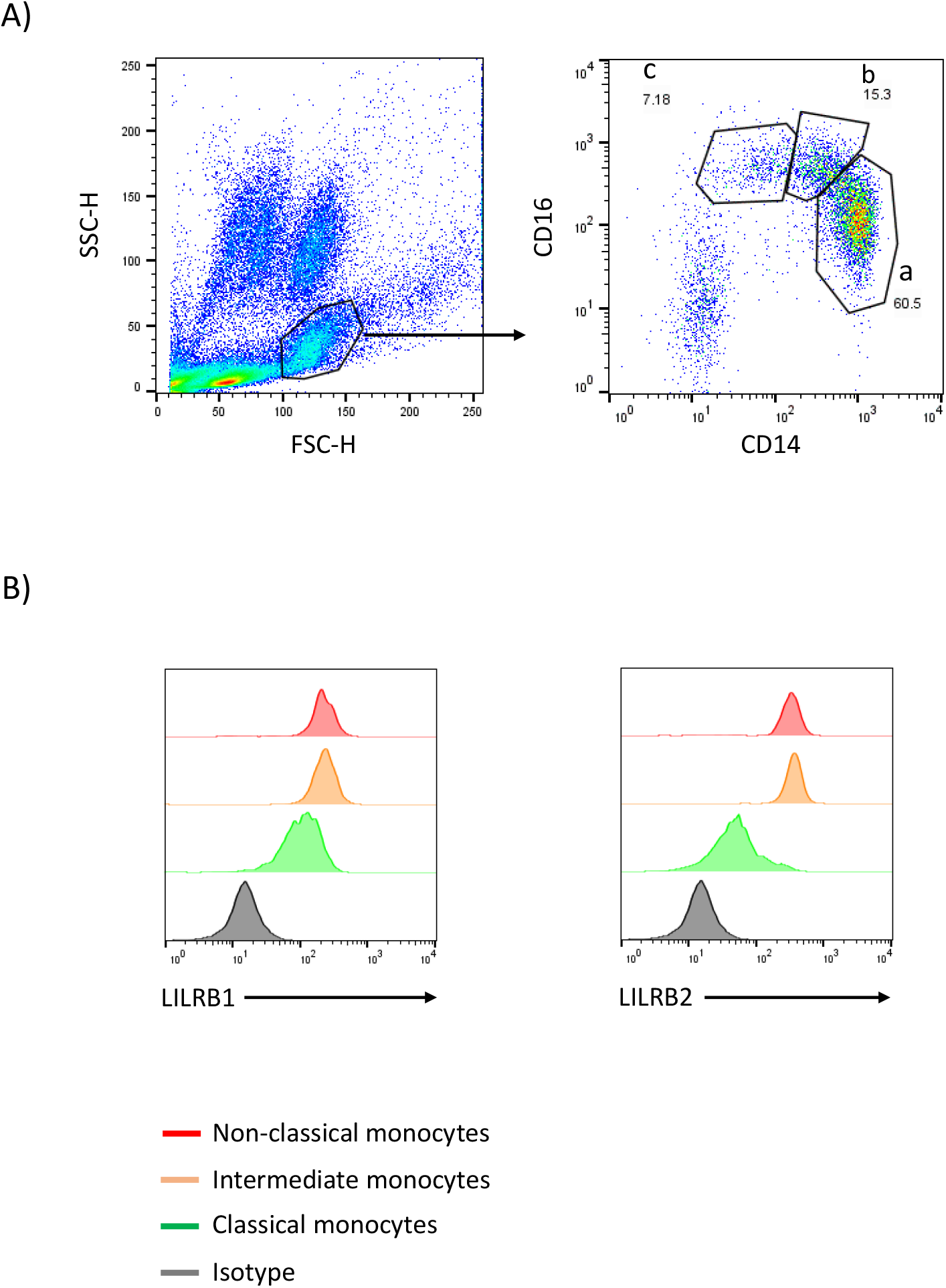
Gating strategy to identify peripheral blood monocyte subsets. Monocytes were gated using forward- and side-scatter properties. The 3 subsets were characterized through expression of CD14 and CD16 markers (a: classical CD14^++^CD16^−^, b: intermediate CD14^++^CD16^+^, and c: non-classical CD14^+^CD16^++^). LILRB1 or LILRB2 expression levels on the 3 monocyte subsets were defined using histogram geometric mean.

### Ethics

The study was conducted in the context of the MiPPAD “Malaria in Pregnancy Preventive Alternative Drugs,” (http://clinicaltrials.gov/ct2/show/NCT00811421) and TOLIMMUNPAL programs. Both projects were approved by the Ethics Committee of the Faculté des Sciences de la Santé de Cotonou. The TOLIMMUNPAL study protocol and informed consent were also approved by the Comité Consultatif de Déontologie et d’Éthique (CCDE) of the Institut de Recherche pour le Développement (IRD, France).

### Statistical analysis

Confounders were environmental exposure (20), infant gender, infant age, birth weight, health centre, ethnic group, maternal anemia and maternal IPTp. A Chi-square test was performed to assess the differences between maternal or neonatal characteristics. The risk of malaria infections between 18 and 24 months of age in infants born to mothers with PM was assessed by a negative binomial regression adjusted on confounders.

Univariate analysis was performed at 18 and 24 months of age using a linear regression to assess (i) the effect of PM on the frequency of immune cell populations and on LILRB1 and LILRB2 expression and (ii) the effect of active malaria infection on LILRB1 and LILRB2 expression, on cytokines and HLA-G levels.

For multivariate analysis, a mixed linear model was used, taking into account the measurement at 18 and 24 months of age together in one analysis. The effect of PM or of an active malaria infection was assessed on the frequency of monocytes, LILRB1 and LILRB2 expression, cytokines, HLA-G and anti-malarial antibody concentrations. All models were adjusted on potential confounders and on the frequency of each subset of immune cell in the LILRB1 and LILRB2 expression analyses. For antibody and cytokines levels analysis, an interaction term was included to determine if the effect of an active malaria infection at the moment of blood draw depends on whether the infant is born to mother with or without PM. Data were analysed with Stata^®^ Software, Version 13 (StatCorp LP, College Station, TX, USA) and the graphs were done using Graph Pad Prism (Version 8.1.2).

## Results

### Infants born to mothers with PM have a higher risk of clinical malaria

The objective of this study was to investigate the immune response at 18 and 24 months of age in infants born to mothers with or without PM. None of the main characteristics of the studied population were found to differ between infants born to mothers with a *Pf*-infected or an uninfected placenta (Table 1). However, infants born to mothers with PM had a significantly higher risk of developing clinical malaria (adjusted negative binomial regression IRR=1.533; [1.020;2.305], p=0.04) between 18 and 24 months of age. There were no differences in the numbers of asymptomatic infections between the two groups of infants.

**Table 1.**
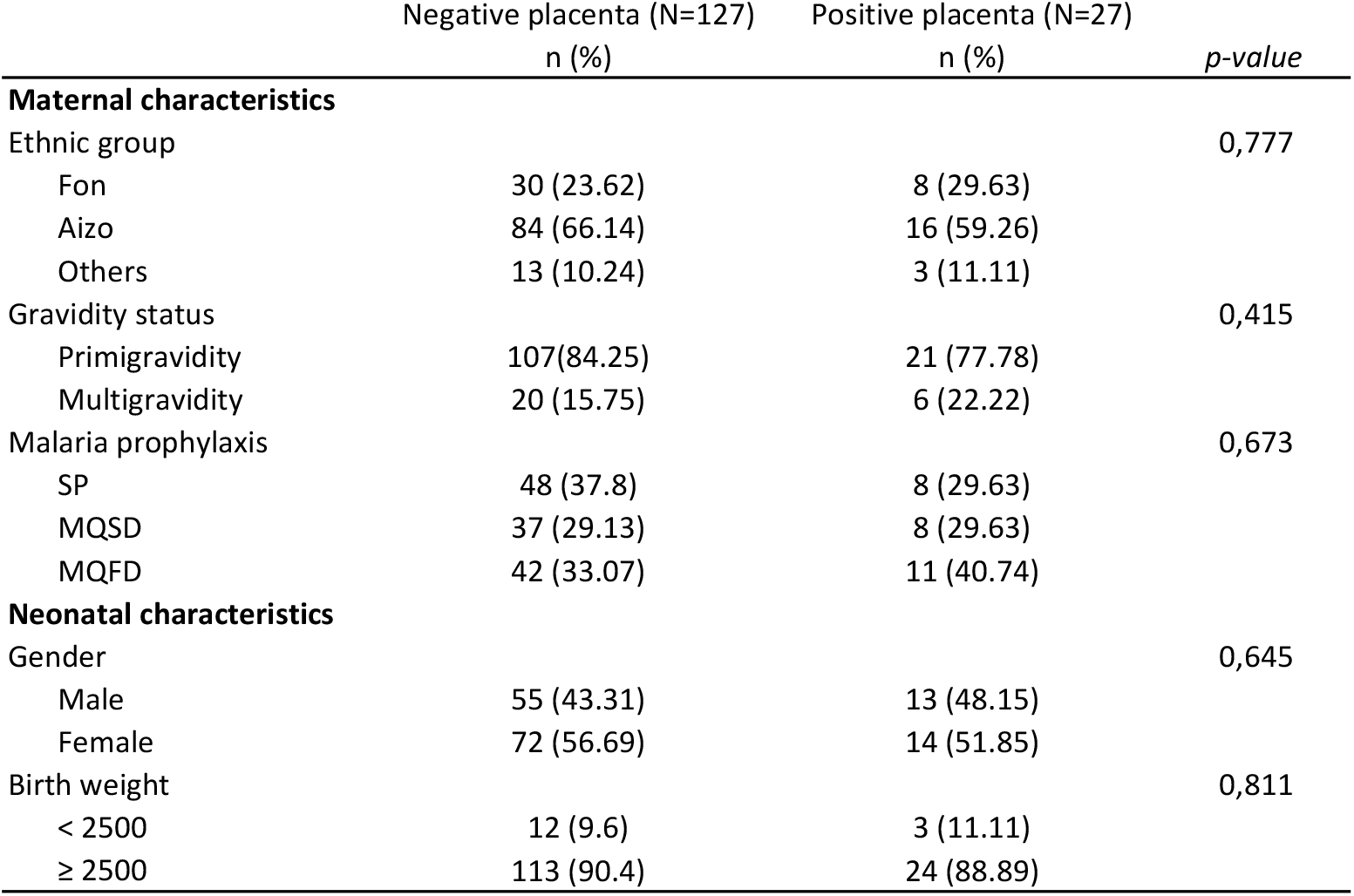
Characteristics of the population. The potential confounding factors are presented. N is the total effective by group (according to PM). Out of 154 newborns, 27 (17.42%) were born from a mother with placental malaria. Chi-square test was performed to compare the confounders between the two groups of infants.

|  | Negative placenta (N=127)<br>n (%) | Positive placenta (N=27)<br>n (%) | *p-value* |
| --- | --- | --- | --- |
| **Maternal characteristics** |  |  |  |
| Ethnic group |  |  | 0,777 |
| Fon | 30 (23.62) | 8 (29.63) |  |
| Aizo | 84 (66.14) | 16 (59.26) |  |
| Others | 13 (10.24) | 3 (11.11) |  |
| Gravidity status |  |  | 0,415 |
| Primigravidity | 107(84.25) | 21 (77.78) |  |
| Multigravidity | 20 (15.75) | 6 (22.22) |  |
| Malaria prophylaxis |  |  | 0,673 |
| SP | 48 (37.8) | 8 (29.63) |  |
| MQSD | 37 (29.13) | 8 (29.63) |  |
| MQFD | 42 (33.07) | 11 (40.74) |  |
| **Neonatal characteristics** |  |  |  |
| Gender |  |  | 0,645 |
| Male | 55 (43.31) | 13 (48.15) |  |
| Female | 72 (56.69) | 14 (51.85) |  |
| Birth weight |  |  | 0,811 |
| < 2500 | 12 (9.6) | 3 (11.11) |  |
| ≥ 2500 | 113 (90.4) | 24 (88.89) |  |

### Non-classical monocytes are less frequent in infants born to mothers with PM and highly express LILRB2

Lymphoid and myeloid immune cells were analyzed in order to assess the effect of PM on infant immunity at 18 and 24 months of age. To this end, a flow cytometry gating strategy was set-up to analyze the three monocyte subsets defined as classical (CD14^++^CD16^-^), intermediate (CD14^++^CD16^+^) and non-classical (CD14^+^CD16^++^) (Figure 1) as well as lymphoid populations, including CD4 T cells, CD8 T cells, NK cells, γδT cells or B cells, neutrophils and eosinophils (Figure S1 and S2). As shown in Figure 2, PM did not alter the proportions of CD4 T cells, CD8 T cells, NK cells, γδT cells, neutrophils, eosinophils or B cells in infants. Interestingly, PM was associated with a higher frequency of classical monocytes and a lower frequency of intermediate and non-classical monocytes. In the multivariate analyses, the proportion of non-classical monocytes remained significantly lower in infants born to mothers with PM, independently of active malaria infection at the time of blood sample collection (Table 2). Of note, malaria infection at the time of sample collection was associated with a reduced frequency of classical monocytes and an increased frequency of intermediate and non-classical monocytes (Table 2). LILRB2 expression was higher on non-classical monocytes, and enhanced to a lesser extent on intermediate monocytes in infants born to mothers with PM as compared to infants from mothers without PM (Table 3). PM did not seem to influence LILRB1 expression by either classical, intermediate or non-classical monocytes (Table 3).

**Figure 2.**
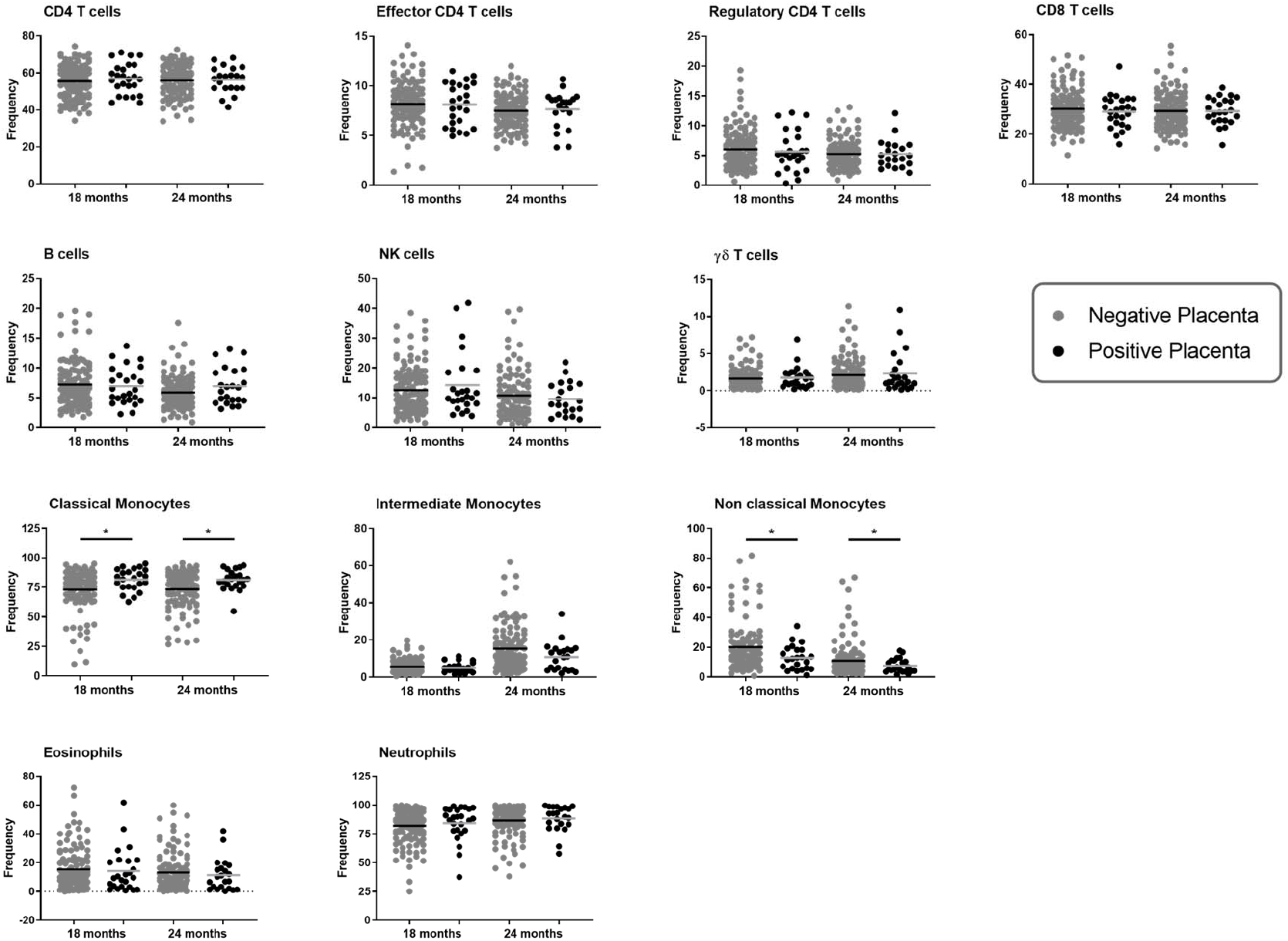
Immune cell populations at 18 and 24 months of age according to placenta malaria. The frequency of each immune cell population at either 18 or 24 months of age was represented in one graphic. Cell subset frequencies were determined as followed: CD4, CD8 and γδ T cells among total CD3^+^ T cells; regulatory and effector CD4^+^ T cells among CD4^+^ CD3^+^ T cells; Monocyte subsets among total monocytes, B or NK cells among lymphoid cells and neutrophils or eosinophils among granulocytes gated using forward- and side-scatter properties. The frequency of immune cells in infants born form mother with or without placental malaria are represented in gray and dark plain round respectively. A linear regression was performed to compare the frequencies between the two groups of infants. *: p value lower than 0.05 (and higher than 0.01).

**Table 2.** Placental malaria decreases non-classical monocyte sub-populations. Adjusted mixed models were used to assess the role of placental malaria on the frequency of monocyte sub-populations at 18 and 24 months of age. Mixed models are used to take into account repetitive measurements for a same individual (dependent variable). For each monocyte sub-population, one model was performed taking into account active *Pf* infection at the sample collection. The model was adjusted on age, gender, birth weight, maternity, ethnicity, maternal anemia, maternal IPTp and environmental exposure. The lincom command (Stata^®^ Software, Version 13 (StatCorp LP, College Station, TX, USA)) was used to compute coefficient values in odds ratios (OR). OR > 1 means that the frequency of monocyte sub-population increases while the frequency of monocyte sub-population decreases if OR<1.

|  | Odds Ratio | p value | 95% Confidence Interval |
| --- | --- | --- | --- |
| Classical monocytes |  |  |  |
| *Pf* infection at sample collect | 0,781 | **<0.001** | [0.711 ; 0.858] |
| *Pf* infection in placenta | 1,146 | **0,013** | [1.029 ; 1.275] |
| Intermediate monocytes |  |  |  |
| *Pf* infection at sample collect | 1,383 | **0,007** | [1.090 ; 1.755] |
| *Pf* infection in placenta | 0,838 | 0,113 | [0.675 ; 1.042] |
| Non classical monocytes |  |  |  |
| *Pf* infection at sample collect | 1,559 | **<0.001** | [0.581 ; 1.999] |
| *Pf* infection in placenta | 0,735 | **0,010** | [0.581 ; 0.930] |

**Table 3.**
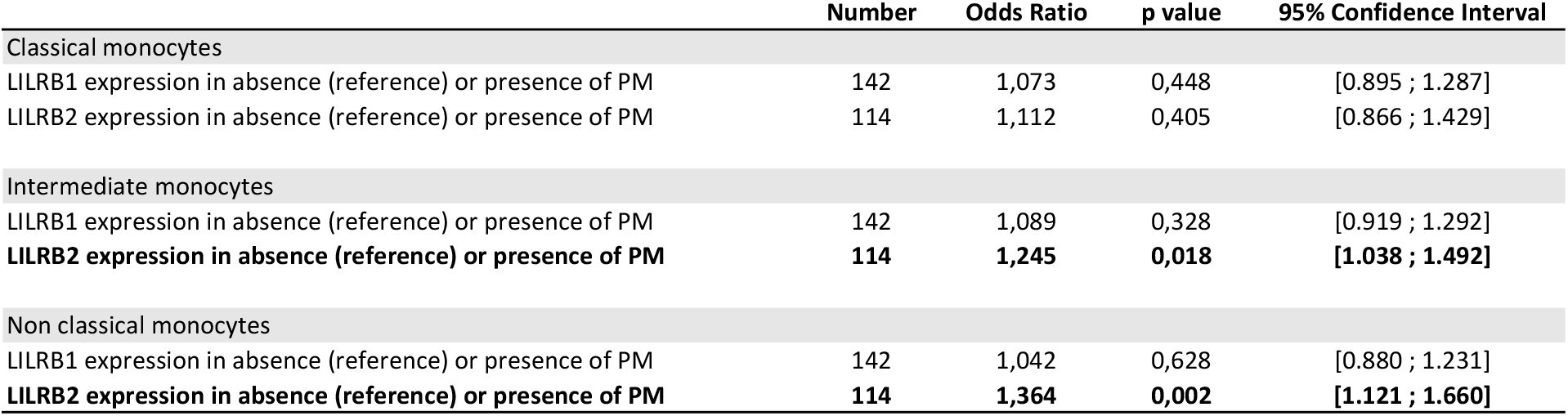
LILRB1 and LILRB2 expression on monocyte sub-populations in infants born to PM-mothers. Adjusted mixed models were used to assess the influence of placental malaria on the LILRB1 and LILRB2 expression on monocyte sub-populations at 18 and 24 months of age. A model was performed for the expression of each inhibitory receptor in the different monocyte sub-populations. The model was adjusted on monocyte subset frequency, age, gender, birth weight, maternity, ethnicity, maternal anemia, maternal IPTp and environmental exposure. Infants with active malaria infection at blood draw were excluded from the analysis. The lincom command (Stata^®^ Software, Version 13 (StatCorp LP, College Station, TX, USA)) was used to compute coefficient values in odds ratios (OR). OR > 1 means that the frequency of monocyte sub-population increases while the frequency of monocyte sub-population decreases if OR<1. A specific analysis was conducted (Figures 3 and 4, Table 4) for infants with active *Pf*-infection.

|  | Number | Odds Ratio | p value | 95% Confidence Interval |
| --- | --- | --- | --- | --- |
| Classical monocytes |  |  |  |  |
| LILRB1 expression in absence (reference) or presence of PM | 142 | 1,073 | 0,448 | [0.895 ; 1.287] |
| LILRB2 expression in absence (reference) or presence of PM | 114 | 1,112 | 0,405 | [0.866 ; 1.429] |
| Intermediate monocytes |  |  |  |  |
| LILRB1 expression in absence (reference) or presence of PM | 142 | 1,089 | 0,328 | [0.919 ; 1.292] |
| **LILRB2 expression in absence (reference) or presence of PM** | **114** | **1,245** | **0,018** | **[1.038 ; 1.492]** |
| Non classical monocytes |  |  |  |  |
| LILRB1 expression in absence (reference) or presence of PM | 142 | 1,042 | 0,628 | [0.880 ; 1.231] |
| **LILRB2 expression in absence (reference) or presence of PM** | **114** | **1,364** | **0,002** | **[1.121 ; 1.660]** |

### Higher LILRB1, LILRB2 and IL-10 expression in infants with active malaria infection

LILRB1 and LILRB2 receptors were expressed at higher levels on intermediate and non-classical monocytes in infants with active *Pf* infection (Figures 3 and 4). In a multivariate analysis, we confirmed that during active *Pf* infection, both receptors were highly expressed on intermediate and non-classical monocytes (Table 4). Moreover, levels of LILRB1 and LILRB2 expression were higher during clinical attacks than during asymptomatic infections (Figures 3 and 4). Finally, out of all the other cell populations evaluated in this project, the levels of LILRB1 were only higher on γδ T cells in infants infected with *Pf* than in infants without malaria infection at blood draw (Figure 5).

**Figure 3.**
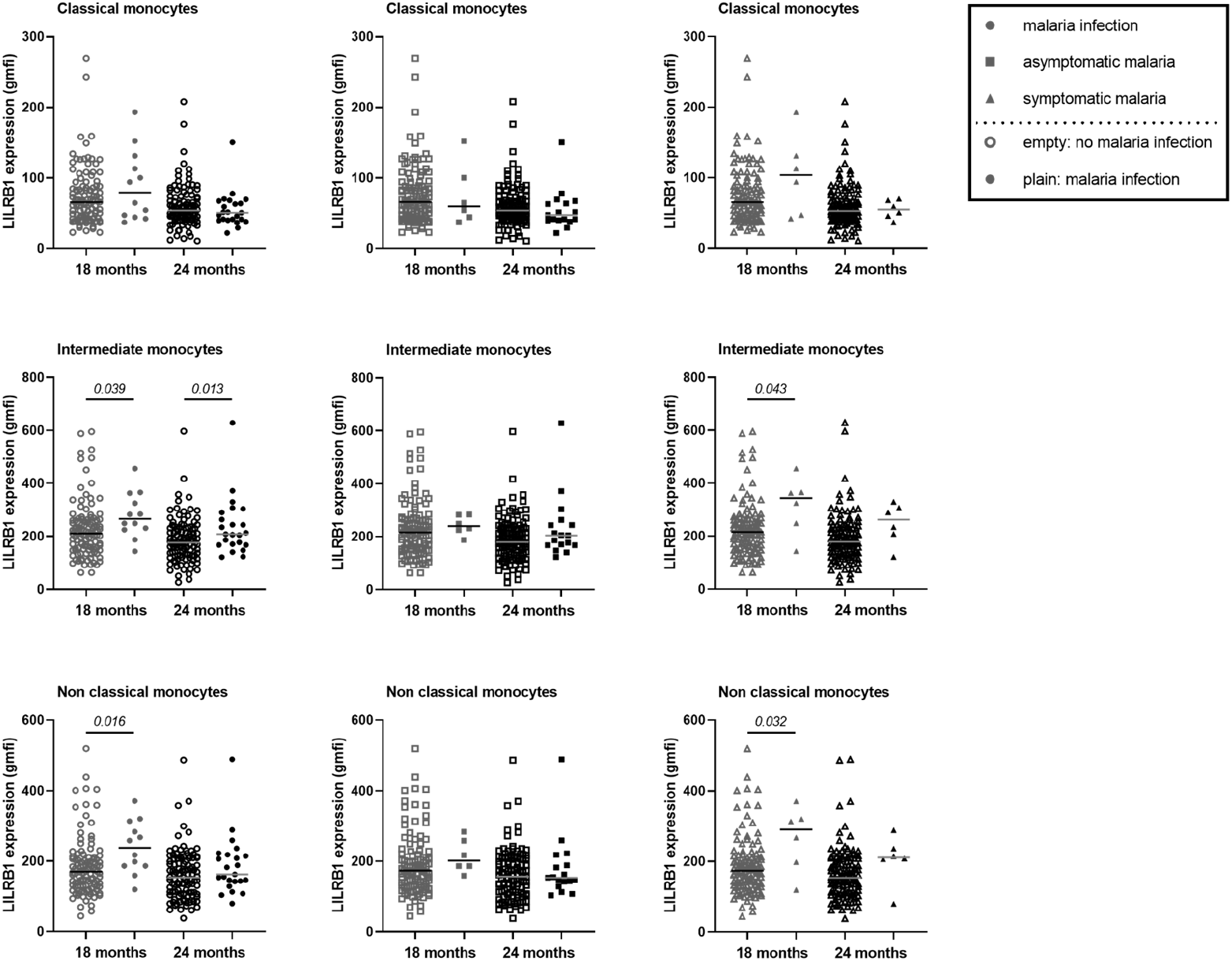
Higher expression of LILRB1 on intermediate and non-classical monocytes at 18 and 24 months of age during active malaria. The expression of LILRB1 on immune cells was measured with the geometric mean of fluorescence intensity (gmfi) between *Pf*-infected and not-infected infants at 18 or 24 months of age. Total, asymptomatic or symptomatic malaria infections were represented with plain circle, square and triangle respectively. No infection at the visit was represented with empty symbols. A linear regression (univariate analysis) was performed to compare the expression between the two groups of infants. P values are indicated in italic.

**Figure 4.**
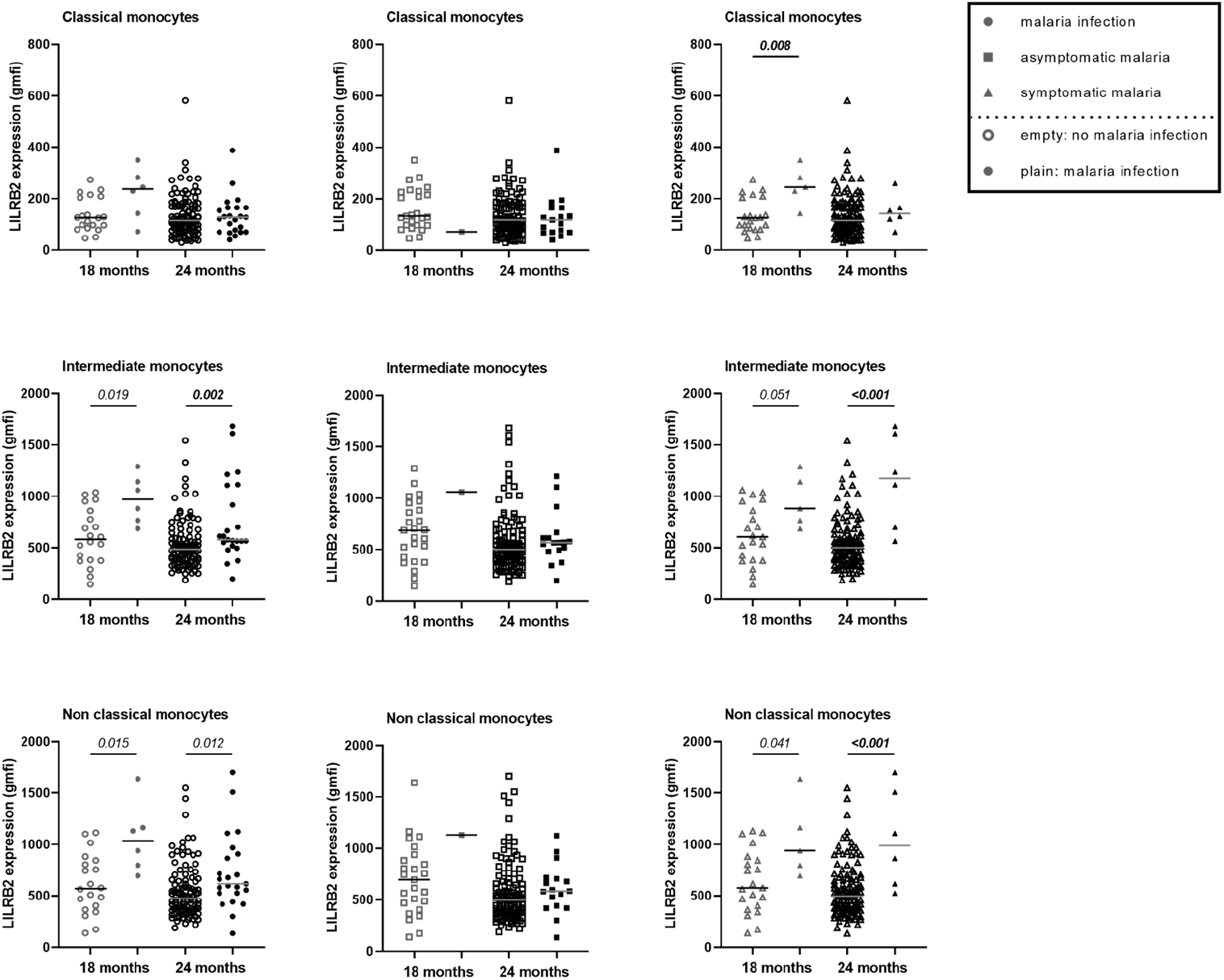
Higher expression of LILRB2 on intermediate and non-classical monocytes at 18 and 24 months during active malaria. The expression of LILRB2 on immune cells was measured with the geometric mean of fluorescence intensity (gmfi) between infected and not-infected infants at 18 or 24 months of age. Total, asymptomatic or symptomatic malaria infections were represented with plain circle, square and triangle respectively. No infection at the visit was represented with empty symbols. A linear regression (univariate analysis) was performed to compare the expression between the two groups of infants. P values are indicated in italic. In bold are the associations still significant after Bonferroni correction.

**Table 4.** Higher expression of LILRB1 and LILRB2 on intermediate and non-classical monocytes at 18 and 24 months during active malaria. Adjusted mixed models were used to assess the influence of *Pf* active infections on the LILRB1 and LILRB2 expression on monocyte sub-populations at 18 and 24 months of age. A model was performed for the expression of each inhibitory receptor in the different monocyte sub-populations. The model was adjusted on monocyte subset frequency, age, gender, birth weight, maternity, ethnicity, maternal anemia, maternal IPTp and environmental exposure. The lincom command (Stata^®^ Software, Version 13 (StatCorp LP, College Station, TX, USA)) was used to compute coefficient values in odds ratios (OR). OR > 1 means that the frequency of monocyte sub-population increases while the frequency of monocyte sub-population decreases if OR<1.

| **Independent variable** | **Number** | **Odds Ratio** | **p value** | **95% Confidence Interval** |
| --- | --- | --- | --- | --- |
| **Classical monocytes** |  |  |  |  |
| LILRB1 expression in absence (reference) or presence of active *Pf* infection | 146 | 1,010 | 0,944 | [ 0.837 ; 1.212] |
| LILRB2 expression in absence (reference) or presence of active *Pf* infection | 150 | 1,051 | 0,671 | [0.820 ; 1.359] |
| **Intermediate monocytes** |  |  |  |  |
| **LILRB1 expression in absence (reference) or presence of active *Pf* infection** | **142** | **1,299** | **0,002** | **[1.102 ; 1.533]** |
| **LILRB2 expression in absence (reference) or presence of active *Pf* infection** | **132** | **1,402** | **<0,001** | **[1.184 ; 1.660]** |
| **Non classical monocytes** |  |  |  |  |
| **LILRB1 expression in absence (reference) or presence of active *Pf* infection** | **146** | **1,279** | **0,002** | **[1.097 ; 1.493]** |
| **LILRB2 expression in absence (reference) or presence of active *Pf* infection** | **132** | **1,362** | **0,001** | **[1.131; 1.640]** |

**Figure 5.**
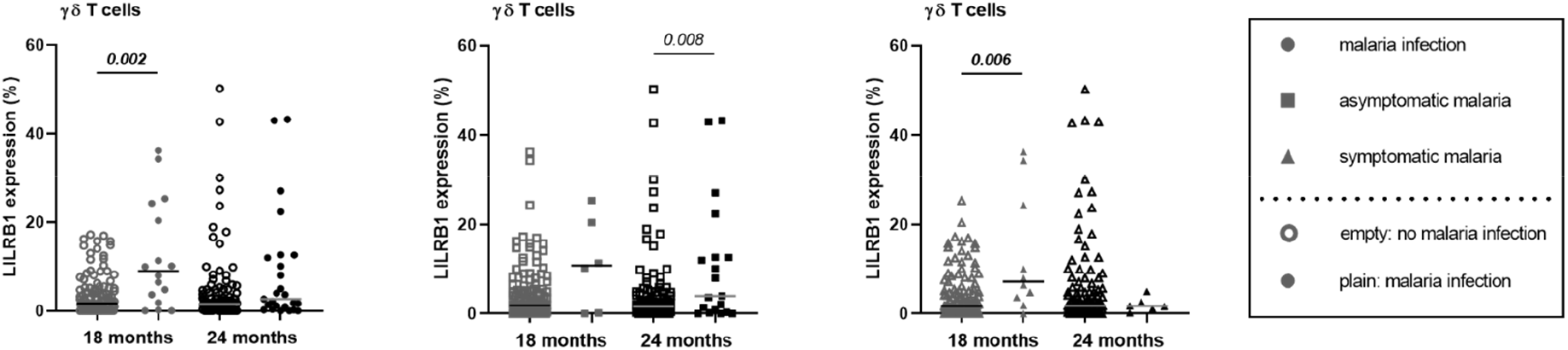
Modulation of LILRB1 expression on γδT cell surface in infant with malaria infections. The expression of LILRB1 on immune cells was measured with the geometric mean of fluorescence (gmfi) in infants at 18 and 24 months of age. Empty circle, square and triangle (gray or black) represented infant without malaria infection at the sample collect. Plain circle, square and triangle (gray or black) represented infant with malaria infection, clinical malaria and asymptomatic infection, respectively. P-values in bold were the one that were still considered significant after Bonferroni correction.

At 18 and 24 months of age, cytokines (IL-4, IFN-γ, TNF-α and IL-10) and soluble HLA-G concentrations were quantified. No differences were observed as a function of PM or of active *Pf* infection except for IL-10 levels. IL-10 levels were higher in those with active malaria infections (febrile and/or asymptomatic (Figure 6)). These observations were confirmed by multivariate mixed models for malaria infections (Coef=2.08; [1.587;2.579]; p<0.001), symptomatic infections (Coef=2.02; [1.455;2.950]; p<0.001) and asymptomatic infections (Coef=1.68; [0.967;2.387]; p<0.001), and mostly in infants with PM (interaction active infection*PM; Coef=2.58; [1.155;4.012]; p<0.001).

**Figure 6.**
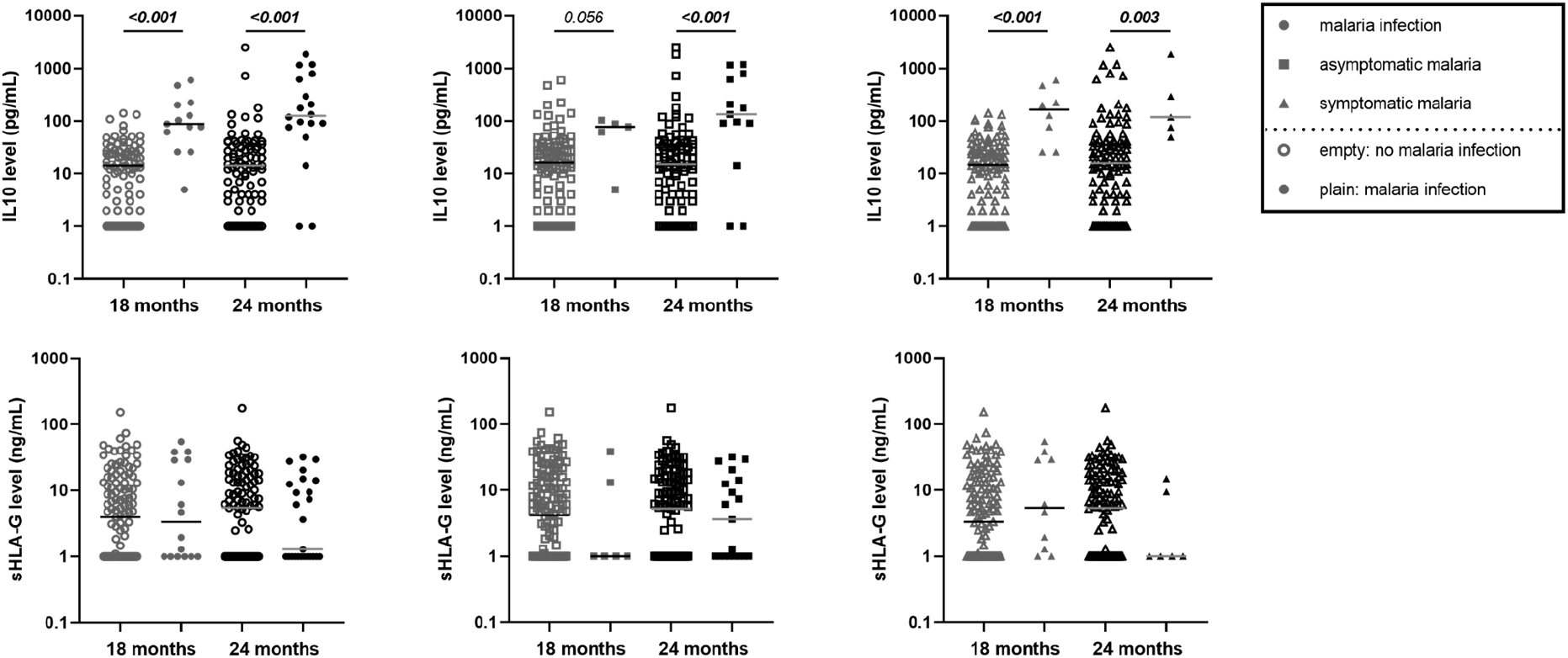
Higher IL10 level in infants with either clinical or asymptomatic malaria infection. The level of IL10 in pg/mL from the plasma of infant at 18 and 24 months of age was represented on the first row of graphics. HLA-G level in ng/mL from the plasma of infant at 18 and 24 months of age was represented on the second row of the graphics. Empty circle, square and triangle (gray or black) represented infants without malaria infection at the time of sample collection. Plain circle, square and triangle (gray or black) represented infant with malaria infection, clinical malaria and asymptomatic infection, respectively. P-values in bold were the one that were still considered significant after Bonferroni correction.

### Lower antibody responses in infants born to mothers with PM

Higher anti-malarial antibody concentrations between 18 and 24 months of age were observed during active malaria infections (Table 5) mostly IgG1 and IgG3 responses against all vaccine candidate antigens (except for IgG3 to MSP1) whereas almost no IgG2 or IgM responses were affected. To understand how anti-malarial antibody levels vary in infants born from PM-mothers during an active *Pf* infection, we introduced an interaction term (interaction active infection*PM, Table 5) in the model. Infants born to mothers with PM seemed less able to mount an antibody responses during active infections (Table 5). Indeed, IgG1 to AMA1, MSP2 and MSP3, IgG2 to MSP2 and IgG3 to AMA1 and MSP2 were significantly lower in *Pf*-infected infants born to PM-mothers (Table 5 and figure 7).

**Table 5.** Lower levels of cytophilic IgG to malaria antigens in infant born to PM-mothers and during active malaria. Multivariate mixed analysis was performed to assess the association between the levels of IgG1, IgG2, IgG3 and IgM specific for seven malaria antigens and active malaria or placental malaria. An interaction term was added to the analysis to test this association in infant born from PM mothers that have an active malaria infection at blood draw. The models were adjusted on age, gender, birth weight, maternity, ethnicity, maternal anemia, maternal IPTp and environmental exposure. PM: Placental malaria, *Pf*: *Plasmodium falciparum*, 95% CI: 95% confidence interval, *: p value<0.05, **: p value<0.01, ***: p value<0.001.

| Antibody level | *Pf* infection |  |  | PM |  |  | Interaction *Pf* infection*PM |  |  |
| --- | --- | --- | --- | --- | --- | --- | --- | --- | --- |
|  | Coefficient | p value | 95% CI | Coefficient | p value | 95% CI | Coefficient | p value | 95% CI |
| **IgG1 to AMA1** | **0.85\*\*** | **0,001** | **[0.338,1.360]** | **0.84\*\*** | **0,003** | **[0.286,1.399]** | -1,15 | 0,050 | [-2.295,0.002] |
| IgG2 to AMA1 | -0,11 | 0,238 | [-0.293,0.073] | **-0.22\*** | **0,043** | **[-0.423,-0.007]** | 0,22 | 0,281 | [-0.184,0.633] |
| **IgG3 to AMA1** | **0.30\*\*\*** | **<0.001** | **[0.162,0.446]** | **0.22\*\*** | **0,005** | **[0.068,0.380]** | **-0.38\*** | **0,019** | **[-0.703,-0.064]** |
| IgM to AMA1 | 0,02 | 0,918 | [-0.359,0.399] | 0,17 | 0,479 | [-0.300,0.640] | -0,48 | 0,393 | [-1.590,0.624] |
| **IgG1 to MSP1** | **1.37\*\*\*** | **<0.001** | **[0.663,2.083]** | -0,22 | 0,571 | [-0.974,0.537] | -1,55 | 0,059 | [-3.157,0.058] |
| IgG2 to MSP1 | -0,01 | 0,938 | [-0.153,0.141] | -0,14 | 0,124 | [-0.311,0.038] | 0,14 | 0,390 | [-0.183,0.469] |
| IgG3 to MSP1 | 0,33 | 0,161 | [-0.129,0.779] | -0,01 | 0,971 | [-0.600,0.578] | 0,01 | 0,990 | [-1.365,1.382] |
| IgM to MSP1 | -0,32 | 0,079 | [-0.676,0.038] | 0,04 | 0,847 | [-0.327,0.399] | 0,50 | 0,231 | [-0.316,1.312] |
| **IgG1 to MSP2-3D7** | **0.58\*** | **0,022** | **[0.085,1.070]** | -0,27 | 0,336 | [-0.819,0.279] | 0,09 | 0,878 | [-1.018,1.191] |
| **IgG2 to MSP2-3D7** | **0.17\*\*\*** | **<0.001** | **[0.083,0.254]** | 0,06 | 0,257 | [-0.047,0.177] | **-0.29\*** | **0,027** | **[-0.556,-0.033]** |
| **IgG3 to MSP2-3D7** | **1.91\*\*\*** | **<0.001** | **[1.111,2.700]** | 0,45 | 0,290 | [-0.384,1.284] | **-2.06\*** | **0,025** | **[-3.864,-0.258]** |
| IgM to MSP2-3D7 | 0,26 | 0,35 | [-0.289,0.816] | 0,32 | 0,270 | [-0.245,0.879] | 0,32 | 0,617 | [-0.939,1.582] |
| **IgG1 to MSP2-FC27** | **0.54\*\*\*** | **<0.001** | **[0.352,0.721]** | **0.21\*** | **0,047** | **[0.003,0.418]** | **-0.79\*\*\*** | **<0.001** | **[-1.201,-0.375]** |
| IgG2 to MSP2-FC27 | 0,03 | 0,379 | [-0.041,0.107] | 0,10 | 0,131 | [-0.030,0.235] | **-0.33\*** | **0,032** | **[-0.633,-0.028]** |
| **IgG3 to MSP2-FC27** | **2.19\*\*\*** | **<0.001** | **[1.532,2.855]** | 0,15 | 0,671 | [-0.554,0.861] | **-1.60\*** | **0,037** | **[-3.091,-0.100]** |
| IgM to MSP2-FC27 | 0,19 | 0,571 | [-0.465,0.843] | 0,33 | 0,328 | [-0.335,1.003] | 0,08 | 0,916 | [-1.412,1.572] |
| **IgG1 to MSP3** | **0.12\*\*** | **0,004** | **[0.038,0.197]** | 0,07 | 0,062 | [-0.004,0.147] | **-0.24\*\*** | **0,006** | **[-0.402,-0.069]** |
| IgG2 to MSP3 | 0,01 | 0,556 | [-0.010,0.018] | 0,01 | 0,472 | [-0.013,0.028] | 0,02 | 0,194 | [-0.010,0.048] |
| **IgG3 to MSP3** | **0.30\*\*\*** | **<0.001** | **[0.162,0.439]** | 0,08 | 0,195 | [-0.043,0.213] | -0,19 | 0,199 | [-0.480,0.100] |
| IgM to MSP3 | -0,03 | 0,482 | [-0.105,0.049] | -0,03 | 0,392 | [-0.112,0.044] | 0,03 | 0,724 | [-0.131,0.188] |
| **IgG1 to GLURP-R0** | **0.25\*** | **0,025** | **[0.031,0.460]** | 0,1 | 0,387 | [-0.128,0.331] | -0,03 | 0,897 | [-0.470,0.411] |
| IgG2 to GLURP-R0 | 0,03 | 0,492 | [-0.058,0.121] | 0,12 | 0,119 | [-0.031,0.273] | 0,04 | 0,704 | [-0.149,0.220] |
| **IgG3 to GLURP-R0** | **0.17\*\*** | **0,003** | **[0.058,0.282]** | 0,01 | 0,992 | [-0.106,0.105] | -0,01 | 0,963 | [-0.240,0.229] |
| IgM to GLURP-R0 | 0,37* | 0,012 | [0.083,0.666] | **0.38\*\*** | **0,007** | **[0.105,0.661]** | -0,46 | 0,138 | [-1.067,0.148] |
| **IgG1 to GLURP-R2** | **0.56\*\*\*** | **0,001** | **[0.245,0.878]** | -0,04 | 0,815 | [-0.331,0.260] | -0,13 | 0,709 | [-0.790,0.537] |
| IgG2 to GLURP-R2 | 0,09 | 0,174 | [-0.039,0.216] | 0,09 | 0,149 | [-0.032,0.213] | 0,11 | 0,408 | [-0.153,0.377] |
| **IgG3 to GLURP-R2** | **0.13\*\*\*** | **0,001** | **[0.056,0.214]** | -0,01 | 0,694 | [-0.084,0.056] | 0,12 | 0,177 | [-0.053,0.285] |
| **IgM to GLURP-R2** | **0.33\*** | **0,039** | **[0.016,0.648]** | 0,02 | 0,903 | [-0.315,0.357] | -0,02 | 0,944 | [-0.674,0.627] |

**Figure 7.**
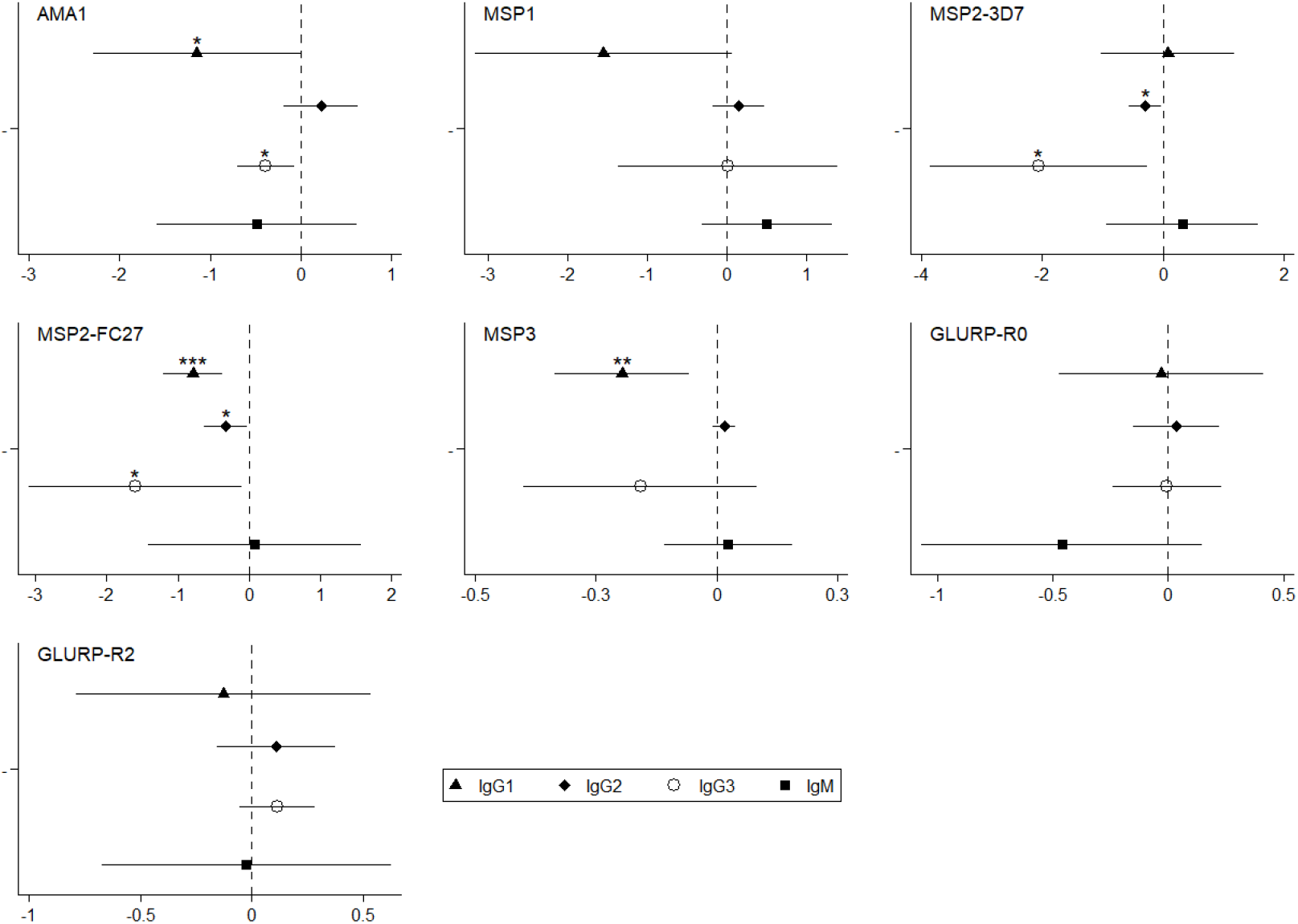
Lower level of specific anti-malarial IgG1 and IgG3 in infant born to PM-mother and with malaria infection. This coefplot showed the synergic effect of PM and *Pf* active infection (interaction term) on the levels of anti-malarial antibodies in infant between 18 and 24 months of age. The models were adjusted on age, gender, birth weight, maternity, ethnicity, maternal anemia, maternal IPTp and environmental exposure.

## DISCUSSION

To our knowledge, this is the first study investigating LILRB1 and LILRB2 inhibitory receptor expression by circulating immune cells in the context of immune-tolerance associated with PM. Our results showed that PM affects monocyte subset frequencies, LILRB2 expression on intermediate and non-classical monocytes, and the levels of cytophilic IgG to *Pf*-merozoite antigens at 18 and 24 months of age.

Monocytes play a crucial role in controlling parasite loads and protecting the host against malaria. Their protective roles include phagocytosis, production of cytokines and antigen presentation. Monocyte-antibody cooperation also generates a major defence mechanism against malaria, known as ADCI (Antibody Dependent Cellular Inhibition) (23), in which monocytes exposed to opsonized merozoites release soluble mediators that inhibit the growth of erythrocytic parasites (24). Higher ADCI levels were associated with clinical protection against malaria (25). Collectively, these studies have emphasized the role of cytophilic IgG1 and IgG3 isotypes in immune mechanisms leading to protection from malaria infection. Their dual ability of opsonizing pathogens and binding tightly to Fc gamma receptors (26) on monocytes or neutrophils designated them as primary actors in the acquisition of natural immunity to malaria (27, 28). On one hand, our study demonstrated that infants born from PM-mothers had lower levels of cytophilic IgG to *Pf*-merozoite antigens between 18 and 24 months of age. On the other hand, we found a lower frequency of non-classical monocytes and a higher level of LILRB2 on those monocytes implying a capacity to modulate (attenuate) cell functions. Taken together, those data suggest impaired opsonizing phagocytic capacity in infants born from PM-mothers. Further studies are needed to confirm this hypothesis of an effect of PM on monocyte functions in infants.

Surprisingly, monocytes that have a relatively short half-life in the peripheral circulation (few days) were affected in 2 years old-infants by maternal PM. Our observations raise questions about the reprogramming of neonatal monocytes/macrophages in PM. Recent studies demonstrate that innate immune cells can build a “memory”. Depending on the nature of the components responsible for the first challenge, monocytes can be trained inducing non-specific attenuated or enhanced innate cell responses and functions after a second challenge (29, 30). This trained memory of human monocytes can persist for at least 3 months (31) and some protective effects have been observed up to 12 months after vaccination with BCG (32). Recent studies have shown metabolic and epigenetic changes in trained innate immune cells (33).. As a result of PM, the fetal immune system can be primed *in utero* (10). Based on our observations and those recent findings (34), we could speculate that a first priming in the fetus could induce a reprogramming of the monocyte precursors reservoir which could then lead to a long-lasting tolerance to infections in infancy. In line with this, Natama *et al*. found that TLR7/8 stimulation of cord blood cells of neonates born to PM-mothers induced higher cytokine levels than that of neonates born to mothers without PM (35). Therefore PM could profoundly alter neonatal immune system, contributing to the higher susceptibility to infections of those infants (35, 36). Further investigations are needed to better characterize the modulation of the neonatal innate immune response (i.e. trained innate immunity) in infants born to PM-mothers.

Another hypothesis to explain the long-lasting effect of PM on infant monocytes could be the capability of various ligands to bind to LILRB1 and LILRB2 and consequently attenuate immune responses. We hypothesized that host molecules known for their anti-inflammatory properties could be involved in such modulation. We measured the plasmatic levels of IL-10 and soluble HLA-G at 18 and 24 months of age. By contrast to previous data in which higher levels of infant HLA-G were found associated with infant malaria infection during follow-up (16), here we observed no association between the levels of HLA-G in infants and active *Pf* infection at time of blood collection. However, we found a higher level of IL-10 during active malaria at both visits and for symptomatic and asymptomatic infections. The concomitant presence of IL-10 and a higher expression of LILRB1 and LILRB2 on non-classical monocytes suggests a down regulation of the cells involved in inflammatory mechanisms and therefore a potentially decreased ability to protect the infants from malaria. Moreover, *in vitro* studies indicate that IL-10 enhances the expression of LILRB2 on monocytes (37). Therefore, the increased expression of LILRB2 on monocytes could result from the production of IL-10 observed in *P. falciparum* infected individuals.

The recently described complex between LILRB1 and the parasite’s family of RIFIN proteins (38) may also be involved in the modulation of monocytes during malaria infections. Indeed, given their strong capacity to attenuate immune response, LILRB1 and LILRB2 represent potential targets for pathogens to evade immune recognition, thereby extending the duration of infection. Indeed, a subset of the RIFINs expressed by *Pf* were shown to bind to LILRB1 and thus lead to the inhibition of B-cell and NK cell functions (39). Our data show an increase of LILRB1 and LILRB2 expression on the non-classical monocyte subsets during malaria infection and in infants born to PM-mothers suggesting that monocytes expressing high levels of LILRB1 and LILRB2 could be more prone to the attenuation of their immune functions possibly through the binding of the RIFINs. The long-lasting tolerance during infancy might be associated with repeated infections involving this particular subset of RIFINs. Further studies are now required to confirm those hypotheses.

Finally, δγT cells play an important role in the immune responses against *Pf* through the production of IFN-γ that is strongly attenuated by LILRB1 engagement (40). Our data show that *Pf*-infection induced an increased frequency of δγT cells expressing LILRB1. Similarly, to monocyte subsets, LILRB1^+^δγT cells could also be inhibited by RIFINs or MHC-I ligands during infection leading to decreased IFN-γ production and cytotoxic activity.

In conclusion, our study provides non-exclusive scenarios for a role of PM in the development of infant immunity to malaria. Interestingly, the role of LILRB inhibitory receptors highlighted in this study raises questions on immune cell regulation pathways and the role of host or pathogen ligands. The impact of such mechanisms would help understand the mechanisms of the natural acquisition of immunity and could therefore contribute to the design of successful vaccine strategies.

## Funding

This study was supported by the Agence Nationale de la Recherche (ANR). Two PhD scholarships were awarded by IRD and the French Embassy, Cotonou to Ibrahim Sadissou and Rafiou Adamou. The clinical trial in which the study was nested was funded by the European Developing Countries Clinical Trials Partnership (EDCTP; IP.2007.31080.002), the Malaria in Pregnancy Consortium. This work was also supported by the Brazil-France research cooperation program USP/COFECUB (grant no. Uc Me 169-17).

## Potential conflicts of interest

The authors declare no conflict of interest.

## Acknowledgements

We would like to thank Georges Snounou and Adrian J.F. Luty for their thorough re-reading of this paper. We are very grateful to the engineers, technicians, midwifes, nurses, drivers, and students from the program and from the two dispensaries where the study took place.

## Notes

### Competing Interest Statement

The authors have declared no competing interest.

