## Supplemental Figures for "LILRB1 and LILRB2 expression in peripheral blood immune cells at 18 and 24 months of age in infants born from mothers with placental malaria"

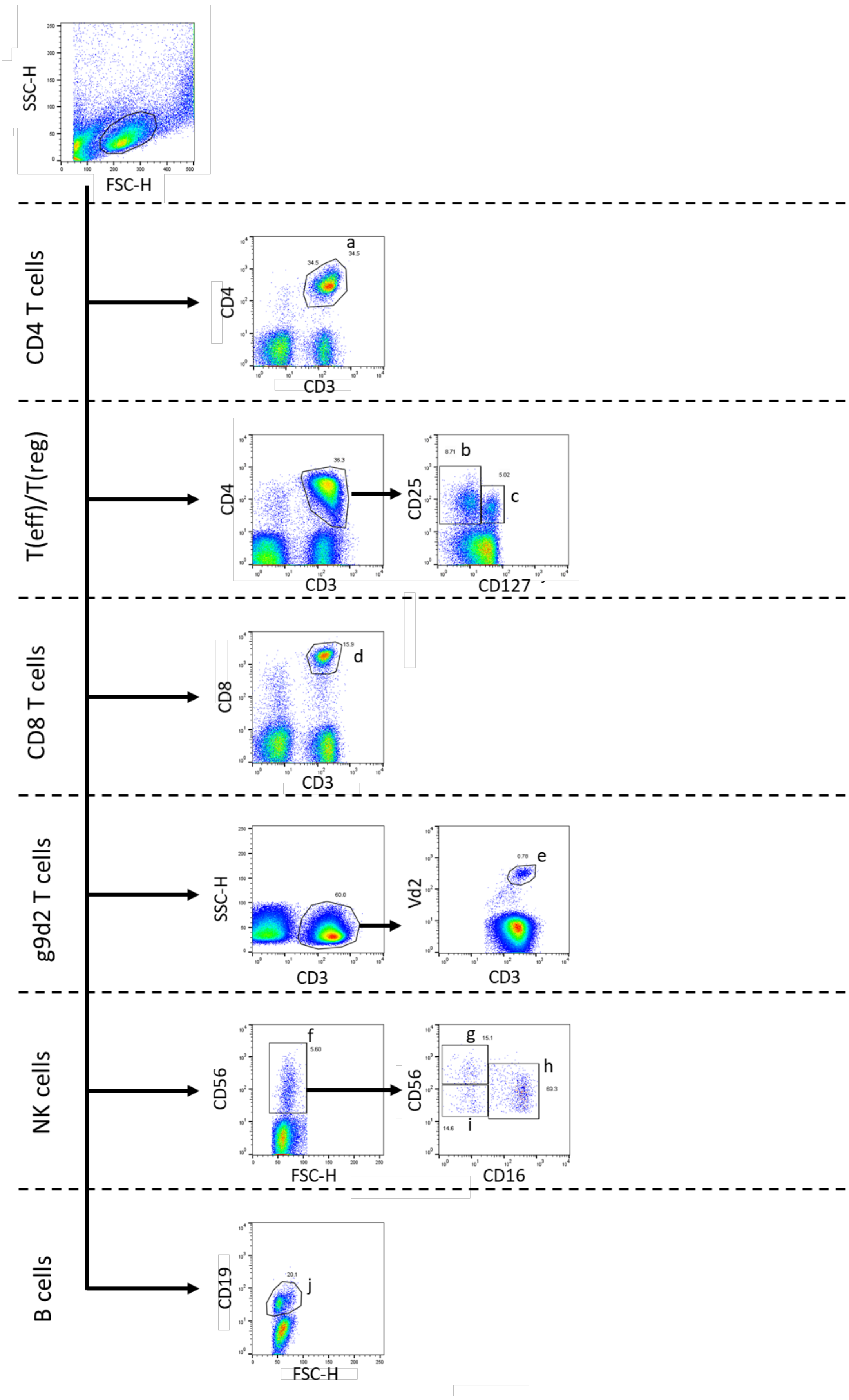


**Figure S1. Gating strategy to define proportions of peripheral blood lymphoid subsets.**

Lymphocytes were gated using forward- and side-scatter properties. Characterization of T helper cells ($CD3^{+}CD4^{+}$), CD8 T cells ($CD3^{+}CD8^{+}$), T regulatory cells ($CD3^{+}CD4^{+}CD25^{high}CD127^{-}$), T effector cells ($CD3^{+}CD4^{+}CD25^{+}CD127^{+}$), $\gamma\delta$ T cells ($CD3^{+}Vd2^{+}$) and B cells ($CD19^{+}$). NK cell subsets were determined based on the expression of CD56 and CD16 markers ($CD56^{mid}CD16^{-}$, $CD56^{dim}CD16^{+}$, $CD56^{high}CD16^{-}$).

a : CD4 T cells ; b : CD4 Treg cells ; c : CD4 Teff cells ; d : CD8 T cells ; e : g9d2 T cells ; f : NK cells ; g : NK cells $CD56^{high}$ $CD16^{-}$ ; h : NK cells $CD56^{dim}$ $CD16^{+}$ ; i : NK cells $CD56^{dim}$ $CD16^{-}$ ; j : B cells

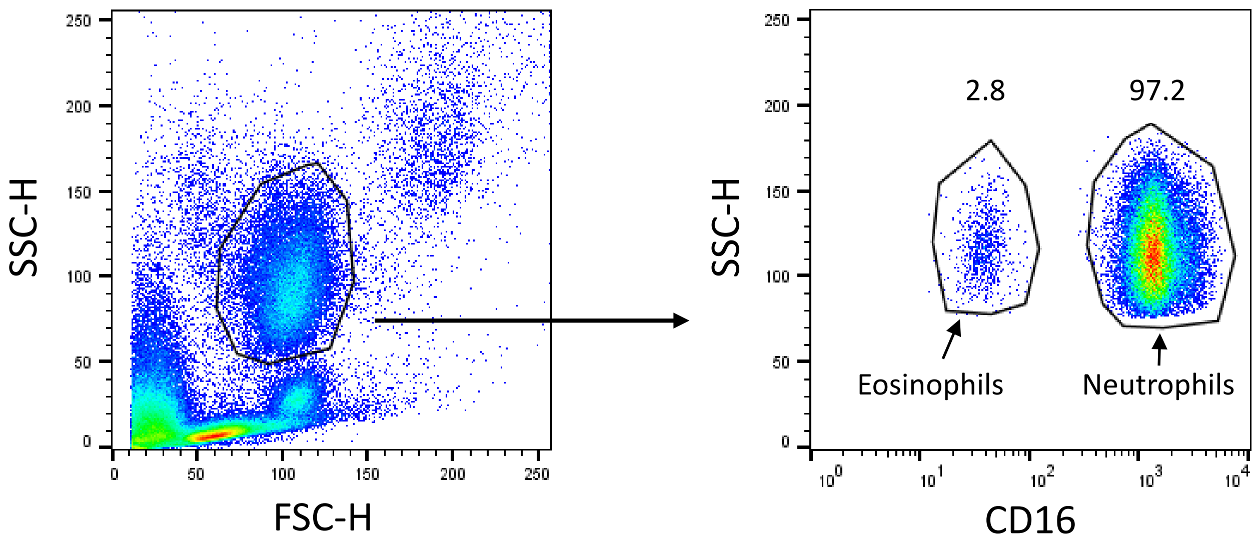


**Figure S2**. **Gating strategy to characterize proportions of peripheral blood neutrophils and eosinophils.**
Granulocytes were gated using forward- and side-scatter properties. Neutrophils were defined as $CD16^{+}$ cells and eosinophils as CD16- cells.
